# Multi-omics profiling reveals atypical sugar utilization and identifies a key membrane composition regulator in *Streptococcus pneumoniae*

**DOI:** 10.1101/2025.06.13.659575

**Authors:** Vincent de Bakker, Xue Liu, Jonah Tang, Matthew Barbisan, Jonathon L. Baker, Jan-Willem Veening

**Author notes:** Correspondence to Jan-Willem Veening.

## Abstract

The human body comprises many different microenvironments, each with their own challenges for microorganisms to overcome in order to survive, and possibly cause infection. The human pathogen *Streptococcus pneumoniae* is notoriously flexible in this regard, and can adapt to a wide range of host niches, including the nasopharynx, lungs, and cerebrospinal fluid. However, the molecular and genetic underpinnings of this ability remain largely obscure. In this work, using infection-mimicking growth conditions we demonstrate that niche adaptation imposes genome-wide changes on multiple levels, including gene essentiality, expression and membrane lipid composition. In general, we show that gene expression and fitness profiling couple orthogonal sets of genes to environmental stimuli. For instance, N-acetylglucosamine (GlcNAc) import (*manLMN*) and catabolism (*nagAB*) genes were required for growth on this sugar, but not differentially expressed in its presence, whereas other amino sugar metabolism pathways were upregulated, but not essential. Surprisingly, we found that pneumococci do not necessarily prefer glucose over GlcNAc and that uptake of GlcNAc in absence of subsequent catabolism was toxic. Moreover, we identified a previously overlooked fatty acid saturation regulator, FasR, controlling membrane composition, rendering it important during heat stress. A fundamental understanding of how genes contribute to bacterial niche adaptation, including nutrient availability or temperature fluctuations, is crucial for understanding successful antibiotic therapy and vaccination strategies and the development of novel anti-infectives.

## Introduction

The human body is the natural habitat of many microorganisms. Various parts of the body impose distinct environmental conditions, such as differences in nutrient availability, surrounding temperature or host immune responses. Nevertheless, some organisms are highly adaptable and manage to survive and grow in multiple human niches. Among these is the bacterium *Streptococcus pneumoniae* (the pneumococcus); a commensal of the human nasopharynx that can cause severe disease states upon invasion of other niches, such as pneumonia in the lungs, or meningitis in the cerebrospinal fluid^1–3^. As such, pneumococci are the main cause of lower respiratory tract infections worldwide, and associated with most deaths in children under five years old^4,5^.

Although it has been well-studied for its pathogenic potential, the genetic foundation enabling its extensive environmental adaptive versatility remains largely uncharted. In fact, as much as 14% of the genes of the highly studied *S. pneumoniae* D39V strain (NCBI accession CP027540) encode proteins of unknown function^6^. Given its apparent need for a high degree of niche flexibility, part of these genes are likely involved in adaptation responses to changing microenvironments. To explore this hypothesis, we have previously quantified gene transcription in several infection-relevant growth conditions by means of chemically defined media^7^. Although insightful, we note that transcriptional changes are generally only moderately reflected on the protein level^8–11^, and not necessarily informative on gene essentiality^12–17^. Indeed, genome-wide Tn-seq coupled to RNA-seq studies in *S. pneumoniae* showed poor links between transcriptional stress responses and gene essentiality^12^.

Here, we address these shortcomings by measuring gene expression on both the transcriptomic and proteomic level, and by interrogating both fitness loss and gain effects of nearly all *S. pneumoniae* D39V genes using CRISPRi-seq^18^. Since this includes baseline essential genes, it allows us to draw comprehensive comparisons between expression levels of all genes and their impact on the bacterium’s fitness, and link these to distinct growth conditions. In this way, we show that niche adaptation happens on all these regulatory levels, and that expression and fitness indeed provide orthogonal, complementary sets of information on the cell state. Specifically, we show that these data can shed light on concrete molecular processes by delving into an unexpected requirement for N-acetylglucosamine catabolism, even in the presence of glucose, and a hitherto overlooked putative membrane composition regulator. Together, these results provide a systems-level perspective of pneumococcal niche adaptation and lay the groundwork for extensive gene function studies.

## Results

### Infection-mimicking growth conditions impose distinct fitness landscapes

Previously, we described pneumococcal transcriptomic profiles for a set of 22 infection-relevant growth conditions, based on chemically defined media^7^. Here, we set out to assess how these differences in expression relate to the importance of those genes for growth in each environment. To this end, we generated genome-wide fitness profiles using CRISPRi-seq with an IPTG-inducible dCas9-sgRNA library in a subset of growth conditions that previously showed clearly distinct transcriptomes^19^.

The selected conditions aim to simulate the host environment as encountered by pneumococci in the nasopharynx (as in colonization), lungs (pneumonia), blood (bacteremia) or cerebrospinal fluid (meningitis), mostly in terms of temperature, acidity, and nutrient availability (Extended Data Table 1)^7^. We also included the commonly used complex media THY and C+Y, the latter both with and without chemically inducing competence by the addition of synthetic competence stimulating peptide 1 (CSP-1), whose transcriptional responses have been well characterized^20–22^.

**Table 1.** FoldSeek relevant top hits with similar folding to predicted SPV_0647 structure. All proteins are transcriptional regulators from the TetR family.

| PDB ID | Bit score | E-value | Probability | Name | Organism | Notes | Reference |
| --- | --- | --- | --- | --- | --- | --- | --- |
| 4MK6 | 233 | $7.54 \cdot 10^{-6}$ | 1.00 | DHSK_reg | *Listeria monocytogenes* | Probable dihydroxyacetone (glycerone) kinase regulator | |
| 5NZ1<br>4DW6<br>5F27<br>5NZ0<br>3Q0V<br>6HO0<br>5NIM<br>5MYM | 187<br>186<br>185<br>185<br>181<br>179<br>175<br>167 | $1.28 \cdot 10^{-4}$<br>$1.55 \cdot 10^{-4}$<br>$4.37 \cdot 10^{-5}$<br>$9.54 \cdot 10^{-5}$<br>$1.05 \cdot 10^{-4}$<br>$1.48 \cdot 10^{-4}$<br>$2.16 \cdot 10^{-4}$<br>$2.41 \cdot 10^{-4}$ | 1.00<br>1.00<br>1.00<br>1.00<br>1.00<br>1.00<br>1.00<br>1.00 | EthR | *Mycobacterium tuberculosis*<sup>48</sup> | Bound and allosterically regulated by medium-chain saturated fatty acid hexadecyl octanoate | Frénois et al. (2004) |
| 6HO6 | 161 | $6.40 \cdot 10^{-4}$ | 1.00 | | | | |
| 2IU5 | 171 | $6.72 \cdot 10^{-4}$ | 1.00 | DhaS | *Lactococcus lactis*<sup>49</sup> | Dihydroxyacetone kinase activator | Christen et al. (2006) |
| 6O6N<br>6O6O | 161<br>158 | $1.09 \cdot 10^{-3}$<br>$1.33 \cdot 10^{-3}$ | 1.00<br>1.00 | FasR | *Mycobacterium tuberculosis*<sup>50</sup> | Activator, binds long fatty acyl-CoA | Lara et al. (2020) |
| 3LSJ | 156 | $1.54 \cdot 10^{-3}$ | 1.00 | DesT | *Pseudomonas aeruginosa*<sup>51</sup> | Binds both unsaturated and saturated fatty acids, regulating their ratio | Zhang et al. (2007) |
| 6OF0 | 155 | $6.40 \cdot 10^{-4}$ | 1.00 | MtrR | *Neisseria gonorrhoeae*<sup>52</sup> | Regulates hydrophobic substrate efflux, hydrophobic binding pocket | Beggs et al. (2019) |
| 5D1W | 153 | $8.57 \cdot 10^{-4}$ | 1.00 | Rv324<br>9c | *Mycobacterium tuberculosis*<sup>53</sup> | Regulates MmpL-mediated lipid export, binds palmitic acid | Delmar et al. (2015) |
| 3WHC | 152 | $1.21 \cdot 10^{-3}$ | 1.00 | FadR | *Bacillus subtilis*<sup>54</sup> | Binds long-chain acyl-CoAs, shown for stearyl-CoA | Fujihashi et al. (2014) |

We observed clear condition-specific fitness landscapes among all conditions tested (Fig. 1a), with 362 operons (corresponding to 615 genes) causing conditional fitness effects (Δ|log2FC|>1, Padj<0.05, Fig. 1b), and a core essentialome of 139 operons (259 genes) (log2FC<1, Padj<0.05, Extended Data Fig. 1a, Supplementary Tables 1-5). We made these essentiality calls easily accessible through our recently updated online genome browser PneumoBrowse 2 (https://veeninglab.com/pneumobrowse)^23^. Apart from the CRISPRi induction effect itself (PC1), the largest source of fitness variation was associated with the difference between complex and defined media (PC2, Fig. 1a). Indeed, most of the genes driving this distinction had metabolic functions, such as members of the Shikimate pathway and the AmiACDEF peptide import system (Fig. 1b, Extended Data Fig. 1b). This was not surprising, since nutrient availability constitutes the biggest difference between the growth conditions used here (Extended Data Table 1). To validate our fitness quantifications, we first sought to confirm such a strong conditional hit, preferably part of a small, well-described pathway with a defined stimulus.

**Figure 1.**
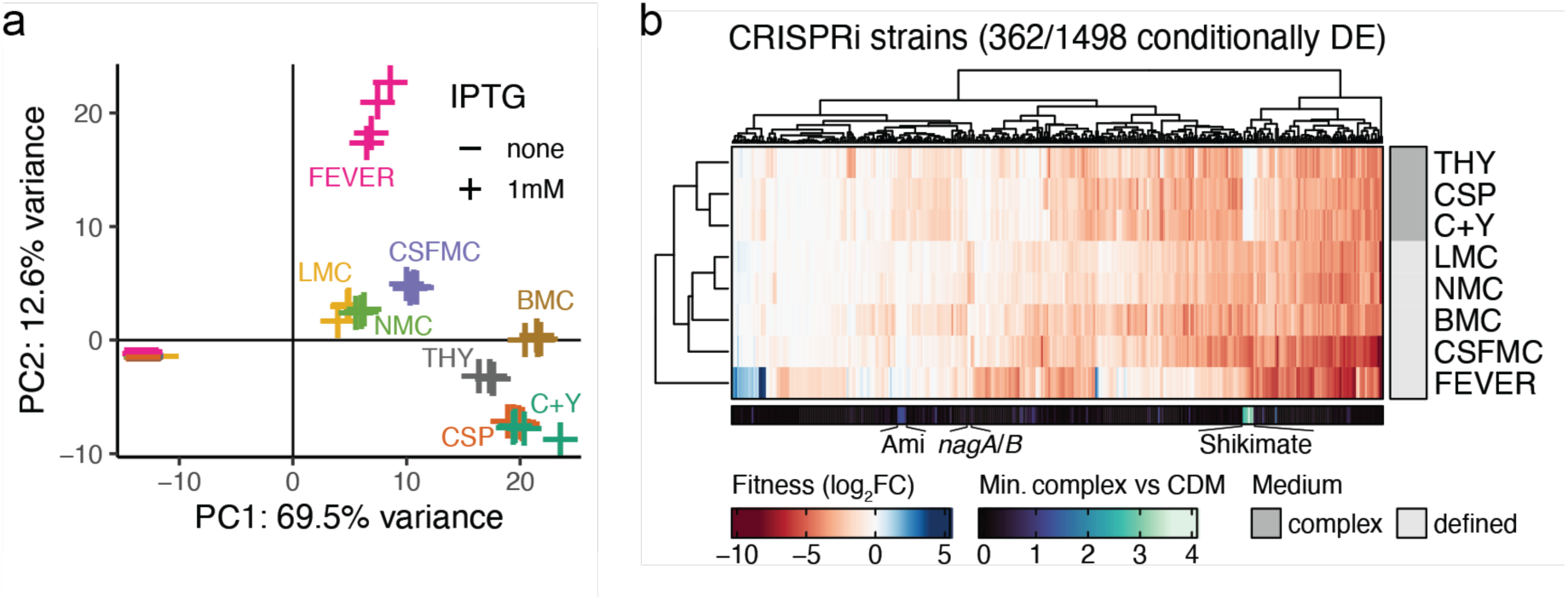
Genome-wide fitness profiles by CRISPRi-seq. (**a**) Principal component analysis demonstrating heterogeneity in CRISPRi strain composition was reproducibly driven by (1) CRISPRi induction following IPTG induction and (2) medium complexity. (**b**) Overview of all CRISPRi strains that were differentially enriched (DE) between at least two growth conditions (Δ|log2FC|>1, Padj<0.05). For each strain, the minimum differential fitness score between any complex versus any chemically defined medium (CDM) condition is also shown, highlighting the strongest consistently differentially essential genes between these medium types. Marked Ami transporter genes: *amiACDEF*; Shikimate genes: *aroACDEK*, *pheA*, *tyrA*.

### N-acetylglucosamine degradation is essential upon import despite glucose availability

The genes *nagA* and *nagB* encode the two proteins responsible for the breakdown of the amino sugar N-acetylglucosamine (GlcNAc), feeding into glycolysis^24,25^. Accordingly, both genes were exclusively essential in the nasopharynx- and lung-mimicking conditions (NMC / LMC), where this was the only sugar added to the medium (Fig. 1B, Extended Data Fig. 1b, Extended Data Table 1). Although we could confirm this result with growth curves of knockout and aTc-inducible complementation strains grown on either GlcNAc or glucose, we also noticed these knockout strains displayed a severe growth defect on an equimolar mix of both sugars (Fig. 2a). This growth defect was reproducible and dose-dependent (Extended Data Fig. 1c). This is surprising, because glucose is commonly believed to be the preferred carbon source in *S. pneumoniae*, which boasts a classic CcpA-based carbon catabolite repression system^26,27^. In addition, neither *nagA* nor *nagB* has any known role in glucose uptake or catabolism. So, in the presence of sufficiently high glucose levels, as in this 1:1 sugar mix, we would expect all strains to grow well, including the mutants.

**Figure 2.**
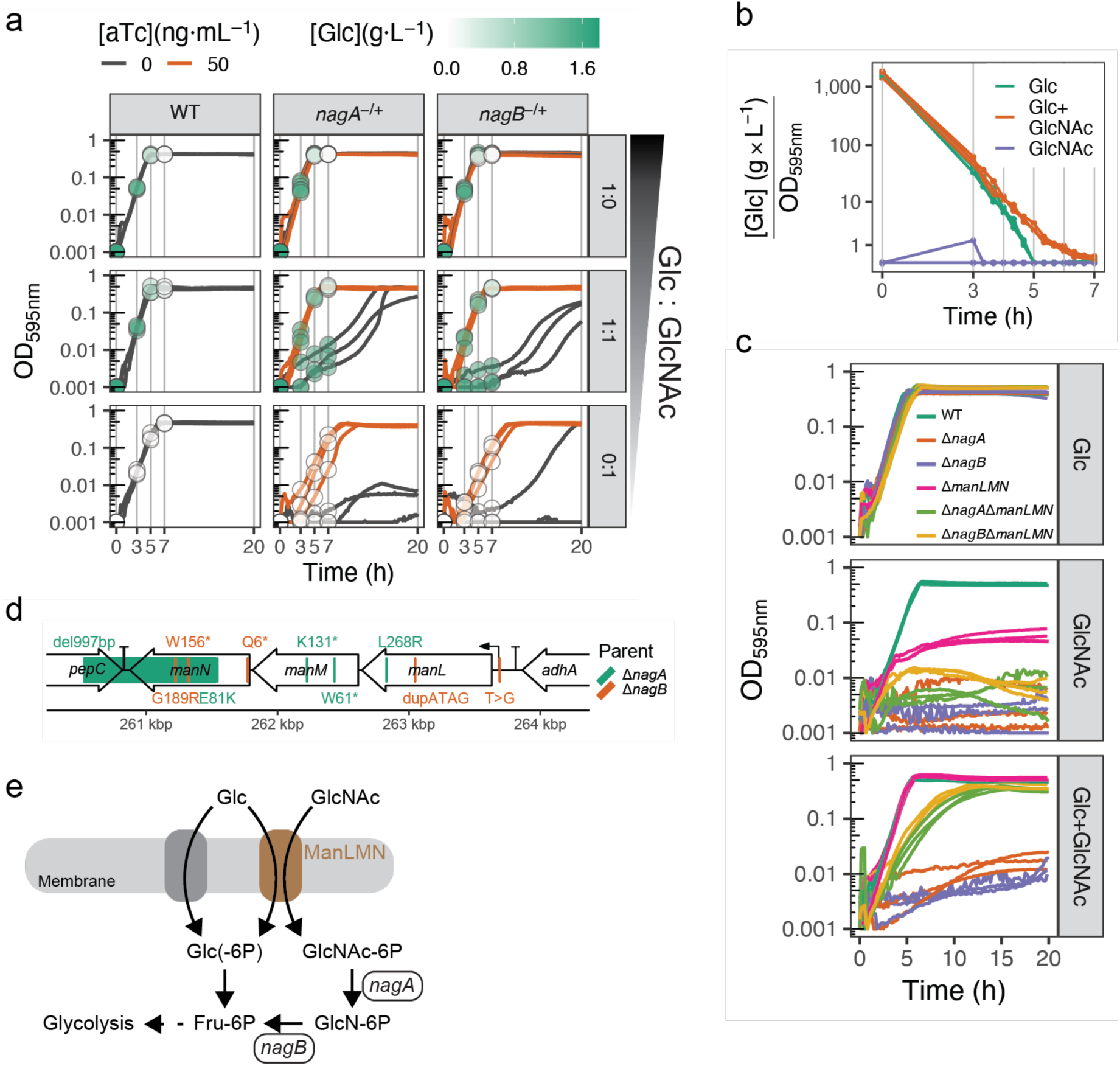
N-acetylglucosamine (GlcNAc) catabolism genes *nagA* and *nagB* are essential on a sugar mix with glucose. (**a**) Growth curves of wild-type (WT) strain as well as aTc-inducible ectopic complementation Ptet-*nagA* and Ptet-*nagB* strains in which the native genes are deleted, respectively (*nagA*^-/+^, *nagB*^-/+^). Glucose (Glc) concentrations in the medium are displayed as points, superimposed on the curves of the replicate they were measured on. Strains were grown with or without aTc on medium with either glucose, GlcNAc, or both in equimolar concentrations. (**b**) Medium glucose concentration normalized by OD for WT. Separate curves for glucose concentration and OD can be found in Extended Data Fig. 1d. (**c**) Growth curves for WT, single and double mutants on medium containing either glucose, GlcNAc, or both in equimolar concentrations. (**d**) Schematic of the *manLMN* locus showing mutations found in *nagA*/*B* suppressor mutants. del: deletion, *: stop codon, dup: duplication. (**e**) Deletion of either *nagA* or *nagB* will result in sugar-phosphate accumulation when grown on GlcNAc, even in the presence of glucose. This accumulation can be alleviated by also deleting *manLMN*, as there are alternative import routes for glucose. Biological triplicates are shown for all experiments.

We therefore decided to measure changes in the glucose concentration in the media during exponential growth using glucose oxidase assays, as a proxy for glucose uptake. Whereas all strains readily depleted the medium of all glucose when no GlcNAc was added, the Δ*nagA* and Δ*nagB* mutants failed to do so in the same time window when both sugars were present (Fig. 2a). Moreover, even in wild-type (WT) bacteria glucose uptake was slower when grown on the sugar mix compared to the glucose-only medium, suggesting GlcNAc presence inhibits glucose import (Fig. 2a). We validated this result by repeating the assay measuring every 20 minutes instead of every two hours, and found a consistent two-hour delay in complete glucose depletion in the presence of GlcNAc (Fig. 2b). Despite this considerable slowdown in glucose uptake, supplementation with GlcNAc did not impact the WT growth rate (Extended Data Fig. 1d). This implies that the bacteria take up GlcNAc even in the presence of glucose, making up for the apparent reduced sugar import to fuel growth. These results suggest that glucose might in fact not necessarily be the preferred carbon source for *S. pneumoniae*.

Since GlcNAc versus glucose uptake seemed competitive, and Δ*nagA* / Δ*nagB* mutants did not appear to import glucose, we hypothesized that their growth defect on the sugar mix was likely caused by the inability to break down GlcNAc after import. Limiting intracellular GlcNAc concentrations by knocking out its importer should then alleviate the growth defect. The main GlcNAc importer has previously been reported to be the phosphotransferase system ManLMN, which was indeed conditionally, albeit not exclusively, essential in NMC and LMC in our CRISPRi-seq assay (Extended Data Fig. 1b)^28,29^. Additionally, *manLMN* deletion caused a strong fitness defect on a GlcNAc-only medium, further confirming its importance in GlcNAc consumption (Fig. 2c). However, on the sugar mix, *manLMN* deletion drastically increased Δ*nagA* and Δ*nagB* viability, in line with our hypothesis (Fig. 2c). Moreover, we performed a suppressor screen by plating both Δ*nagA* and Δ*nagB* mutants on agar containing both sugars. We isolated 20 colonies, 10 for each parent strain, all of which phenocopied the partial rescue of the double knockout strains in liquid (Extended Data Fig. 1e). Sequencing results of 10 of these isolates, five for each parent strain, revealed that all of them were mutated in the *manLMN* locus, indeed suggesting selection for GlcNAc import reduction (Fig. 2d).

Taken together, our results reveal an unusual sugar preference in *S. pneumoniae* and imply toxic intermediates in the GlcNAc catabolic pathway. Since all intermediates are sugar phosphates (Fig. 2e), and the toxicity is seen in both the Δ*nagA* and Δ*nagB* mutants (Fig. 2b,d), we speculate this could be a form of sugar-phosphate stress, as has been described for other species and sugars^30^.

### Heat stress imposes specific genetic requirements

Although metabolic effects dominated the conditional fitness profiles by separating complex media from defined ones, the FEVER condition appeared more distinct (Fig. 1a-b). Since bacteria in this condition were consistently grown at an elevated temperature of 40°C, we sought to explore whether this could explain FEVER-specific effects. To this end, we compared this growth condition with CSFMC, where the ambient temperature was 37°C, but which was otherwise identical (Extended Data Table 1).

Strikingly, we uniquely observed fitness gain effects for the knockdown of certain genes in FEVER (Fig. 1b). Among the strongest of these was arginine metabolism regulator *argR1* (Fig. 3a). ArgR1 functions as a dimer with AhrC, whose gene indeed had a similar effect on fitness^31^. Since these two genes are located more than 780 kbp apart, this cannot be due to CRISPRi polar effects. Most other strong fitness gains were achieved through repression of genes neighboring *ahrC* (*xseA*, *xseB*, *ipsA*-*spv_1064*, *recN*), which we therefore do expect to be due to polarity (Fig. 3a, Supplementary Tables 4-5). In addition, arginine is the precursor for polyamine biosynthesis in *S. pneumoniae*, and most of the genes involved in this pathway also showed mild fitness gain effects upon repression, of which four significantly so (log2FC>1, Padj<0.05, Fig. 3a, Supplementary Tables 4-5)^32^. Knockdown of these genes thus likely increases intracellular arginine levels, an effect also associated with *ahrC* and *argR1* repression^31^. Together, these results suggest arginine retention is favorable for *S. pneumoniae* at higher temperatures, although this remains to be investigated more deeply.

**Figure 3.**
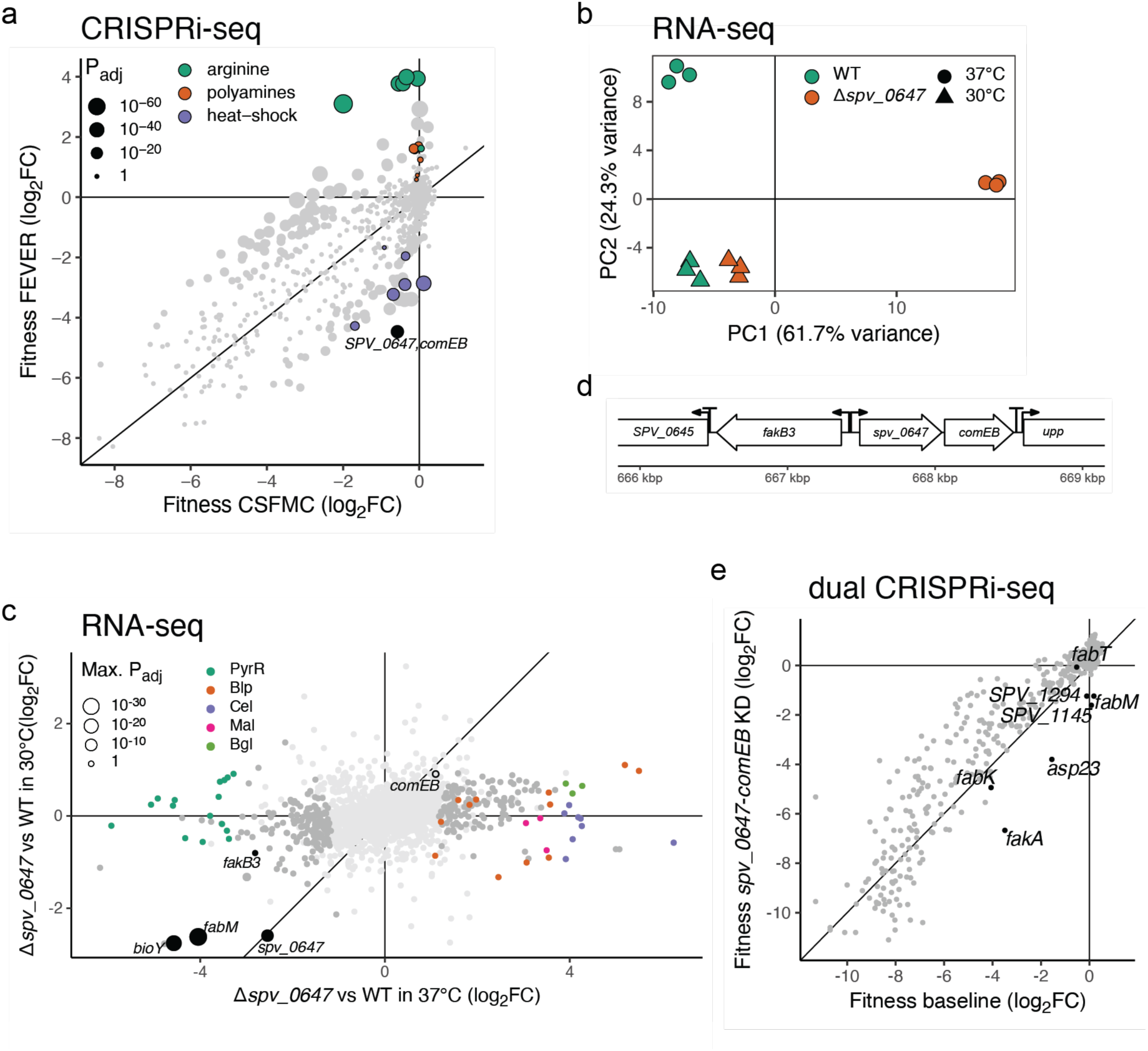
*spv_0647* is important during heat stress and functionally related to genes modulating fatty acid saturation levels. (**a**) Fitness in FEVER condition (40°C) compared to CSFMC (37°C) as measured by CRISPRi-seq. Adjusted P-values represent significance testing for a fitness difference of at least one log2FC. (b) Principal component analysis of Δ*spv_0647* mutant and WT transcriptomes measured by RNA-seq at either 30°C or 37°C; based on rlog normalization^34^. (**c**) Differential expression of genes between the Δ*spv_0647* mutant and WT strains at either 37°C or 30°C. The maximum adjusted P-value representing differential expression at either temperature is shown per gene. Genes not differentially expressed at either temperature (Padj>0.05 for |log2FC|>1) are displayed in light grey. (**d**) Genomic neighborhood of *spv_0647*, including promoters and terminators. (**e**) Genetic interactions of *spv_0647* according to dual CRISPRi-seq data of Dénéréaz *et al.* (2024)^35^. Shown are the fitness effects of each sgRNA in the absence (baseline) or presence of the sgRNA targeting the *spv_0647*-*comEB* operon.

As expected, repression of genes encoding known heat-shock proteins such as *groEL*/*ES*, *ftsH* and the *hrcA*-*grpE*-*dnaK*-*spv_2171*-*dnaJ* operon resulted in strong fitness losses (Fig. 3a, Supplementary Tables 4-5)^33^. The strongest FEVER-specific fitness loss was however caused by a two-gene operon hitherto not associated with heat stress: *comEB*-*spv_0647* (Fig. 3a). We first validated this result with an operon-based deletion and complementation strain, and subsequently confirmed that deletion of *spv_0647*, but not *comEB*, causes growth defects at higher temperatures (Extended Data Fig. 2a).

### SPV_0647 is critical for membrane homeostasis during high temperatures

Since *spv_0647* encodes a hypothetical, putative transcriptional regulator (TetR family), we made a clean knockout strain and performed RNA-seq comparing its transcriptome to that of WT at two different temperatures, in order to uncover its potential regulon (Supplementary Tables 6-9). To limit toxic side effects, and because the growth defect was already visible at 37°C, we opted to draw the comparison between this temperature and 30°C, where the growth defect of the mutant was virtually absent (Extended Data Fig. 2a).

Indeed, the transcriptomes diverged substantially at 37°C, confirming a temperature-dependent effect (Fig. 3b). While many more genes were differentially expressed at the higher temperature, *spv_0647* mRNA was always depleted in the mutant, as expected (Fig. 3c). Similarly, multiple genes involved in fatty acid metabolism were also consistently downregulated, most notably *fabM*, *fakB3* and fatty acid biosynthesis cofactor biotin transporter *bioY* (Fig. 3c).

We argued that dysregulation of fatty acid metabolism could alter membrane composition, affecting properties such as its permeability and fluidity. As temperature changes themselves are also known to affect these properties, we hypothesized that the growth defect of the mutant at higher temperatures is the result of an interaction between heat stress and fatty acid dysregulation, compromising membrane integrity. In turn, membrane weakening would likely disrupt many downstream processes, especially those involving transmembrane transport or membrane-anchored proteins. This could explain the massive, global transcriptome divergence we observed at 37°C, including pyrimidine biosynthesis (PyrR), bacteriocin production (Blp) and multiple sugar metabolism (maltose, (hemi)cellulose, beta-glucoside) loci (Fig. 3c).

Fatty acid biosynthesis in *S. pneumoniae* is controlled by FabT, regulating expression of the biosynthesis FASII operon^36^. However, the first gene of the locus, *fabM*, is not controlled by FabT^37^. FabM balances substrate availability for the production of unsaturated versus saturated fatty acids, and a mutant cannot synthesize unsaturated variants^38,39^. Although pneumococci can only make mono-unsaturated fatty acids, they are able to take up exogenous poly-unsaturated fatty acids through binding of FakB3 and subsequent phosphorylation by FakA, making these species also available for membrane incorporation^40–44^. As such, downregulation of either *fabM* or *fakB3* should yield relatively higher saturated to unsaturated fatty acid ratios (SFA:UFA) in the membrane, which is indeed known to be a major factor affecting membrane properties like fluidity^45,46^. Moreover, *fakB3* is located directly upstream of *spv_0647* on the chromosome, in antisense orientation (Fig. 3d). Since transcriptional regulator genes are often situated adjacent to the genes they regulate, it is tempting to speculate on a potential *fakB3*-regulating role for SPV_0647.

To get more clues regarding SPV_0647 function, we compared its predicted folding to determined crystal structures in PDB using FoldSeek^47^. Strikingly, we found that despite strong sequence dissimilarities, the protein is predicted to have a structure similar to multiple known TetR-like lipid metabolism transcriptional regulators, including activators (Table 1, Extended Data Fig. 2b, Supplementary Table 10). These hits point to a role of SPV_0647 in fatty acid regulation.

We next revisited our recently published *S. pneumoniae* D39V genome-wide genetic interaction data generated by dual CRISPRi-seq^35^, and found negative interactions of the *spv_0647*-*comEB* operon with *fakA* and *fabM* (Fig. 3e). Strikingly, this effect was not observed for the other two sgRNAs targeting the directly downstream *fabT* and *fabK* in the FASII locus. This implies the interaction only involves *fabM*. Furthermore, *asp23* displayed a similar negative interaction, which is likely a polar effect, given its location directly upstream of *fakA* without intermediate terminator^6^. Furthermore, the negative interaction with *spv_1145*, a dNTP triphosphohydrolase, is likely due to the polar knockdown of *comEB*, a dCMP deaminase. Lastly, the genetic interaction with *spv_1294* implies a potential role for the encoded hypothetical protein in either DNA or fatty acid metabolism. These results suggest *spv_0647* might have a function in FakA- and FabM-related processes, influencing fatty acid saturation levels in the membrane.

### FasR (SPV_0647) controls fatty acid saturation balance in the membrane mediated by *fabM*

To test whether the growth phenotype was brought about by *fabM* or *fakB3* downregulation (Fig. 3c), we tried to rescue the bacteria by overexpression of either or both genes in the Δ*spv_0647* mutant. *fakB3* overexpression improved growth marginally at best, which was not surprising, as the growth medium did not contain an explicit excess of poly-unsaturated acids. However, *fabM* overexpression clearly restored growth (Fig. 4a). This implies that generating a surplus of (substrate for) unsaturated fatty acids rescues the mutant, which might thus lack sufficient levels of these fatty acid species.

**Figure 4.**
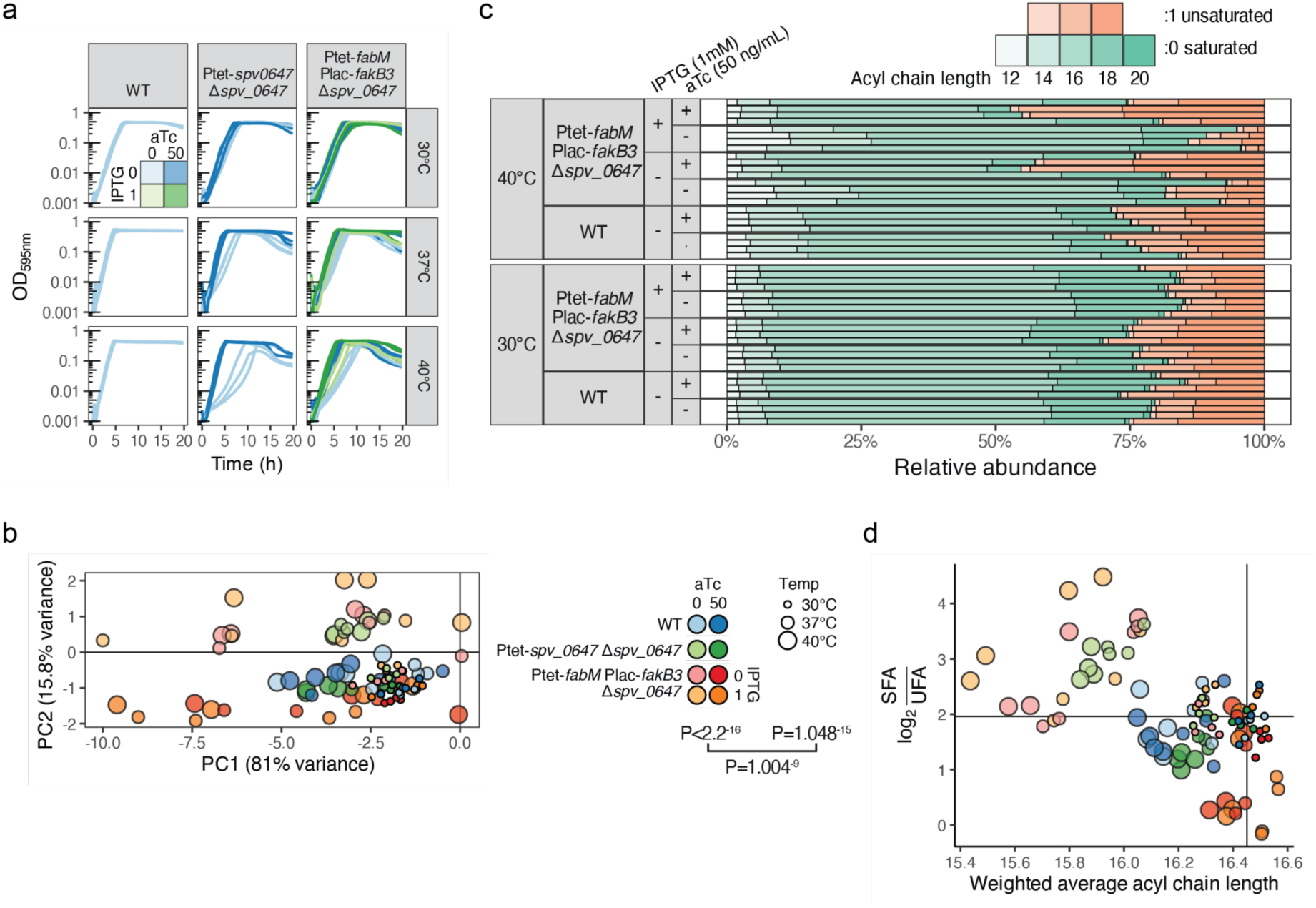
*fasR* affects heat resistance by modulating membrane composition. (**a**) Growth curves of Δ*spv_0647* with either aTc-inducible complementation or aTc- and IPTG-inducible *fabM* and *fakB3* overexpression, respectively, at different temperatures. Biological triplicates are shown. (**b**) Principal component analysis of membrane composition profiles (as in Extended Data Fig. 2c), based on Aitchison composition (acomp) transformation^55^. P-values of temperature, strain- induction, and their interaction effects on fatty acid composition are shown (compositional ANOVA, irl-transformed^55^). (**c**) Relative fatty acid composition in membranes of selected strains and temperatures, measured by GC-FAME. The complete set of samples can be found in Extended Data Fig. 2c. (**d**) Average acyl chain length per sample, weighted by relative fatty acid abundance, versus the ratio between summed relative levels of saturated and unsaturated fatty acids. Vertical and horizontal lines represent averages for WT grown without aTc at 30°C, for reference. Legend is shared with panel b.

Finally, we aimed to assess such changes in the composition of the membrane itself. Using gas chromatography-mass spectrometry of fatty acid methyl esters (GC-FAME), we detected eight fatty acid species in the membranes of WT, mutant, complementation and overexpression strains grown to mid-exponential phase at 30°C, 37°C and 40°C (Extended Data Fig. 2c, Supplementary Table 11), and found that temperature affected the obtained composition profiles in a strain-dependent fashion (compositional ANOVA P<0.05, Fig. 4b). Indeed, they were similar at 30°C but diverged at higher temperatures (Fig. 4b-c), matching the growth and transcriptome phenotypes observed before (Extended Data Fig. 2a, Fig. 3b). Temperature increase was generally associated with a slight decrease in average acyl chain length, but specifically in the Δ*spv_0647* mutant with an inability to maintain unsaturated fatty acid levels (Fig. 4c-d), in line with *fabM* downregulation (Fig. 3c). This phenotype could not only be rescued by *spv_0647* complementation, but also by *fabM* overexpression, which could even cause an overproduction of unsaturated fatty acids (Fig. 4c-d, Extended Data Fig. 2c). These results correspond exactly to the growth phenotypes we observed for the same strains (Fig. 4a).

Together, these findings support a model in which SPV_0647 confers heat resistance by modulating the SFA:UFA balance in the membrane through positive regulation of *fabM* and possibly *fakB3*. Therefore, we have renamed SPV_0647 to FasR, for fatty <u>a</u>cid <u>s</u>aturation regulator.

### Differences in gene expression do not reflect differences in fitness effects

We next wanted to know to what extent these and other differential fitness requirements translate to expression patterns. To this effect, we measured both the transcriptome and proteome of *S. pneumoniae* D39V WT grown in six of the infection-mimicking conditions in which CRISPRi-seq was performed (Extended Data Table 1). Using RNA-seq and quantitative, label-free LC-MS, we detected 2146 different RNA species and 870 proteins, of which we retained 2140 and 736, respectively, following standard normalization and imputation methods (Supplementary Tables 12-17). Small, lowly abundant, and membrane proteins were underrepresented in the proteomics measurements (Extended Data Fig. 3a-c). Normalized quantifications of these data were made easily accessible in PneumoBrowse 2^23^. In general, transcriptome-proteome correlations resembled those reported before for other organisms^8,11^. Briefly, transcript and protein levels correlated moderately (R^2^ = 0.37-0.51) within growth conditions (Extended Data Fig. 3d), and protein levels tended to increase with transcript levels of individual genes across conditions (Extended Data Fig. 3e). Intuitively, this effect was strongest for genes of which both transcript and protein products were differentially enriched between at least two growth conditions (Extended Data Fig. 3f), indicating tighter mRNA-protein regulation.

Moreover, dimension reduction by Multi-Omics Factor Analysis (MOFA)^56^ showed reproducible, integrated, condition-specific proteo-transcriptomic landscapes, while accounting for both the paired nature of the data and non-coding transcripts (Fig. 5a). Its first two dimensions (factors) explained most of the variance in either data set (Extended Data Fig. 4a), correlated well with the most highly differentially expressed genes (Fig. 5b), and revealed three main patterns.

**Figure 5.**
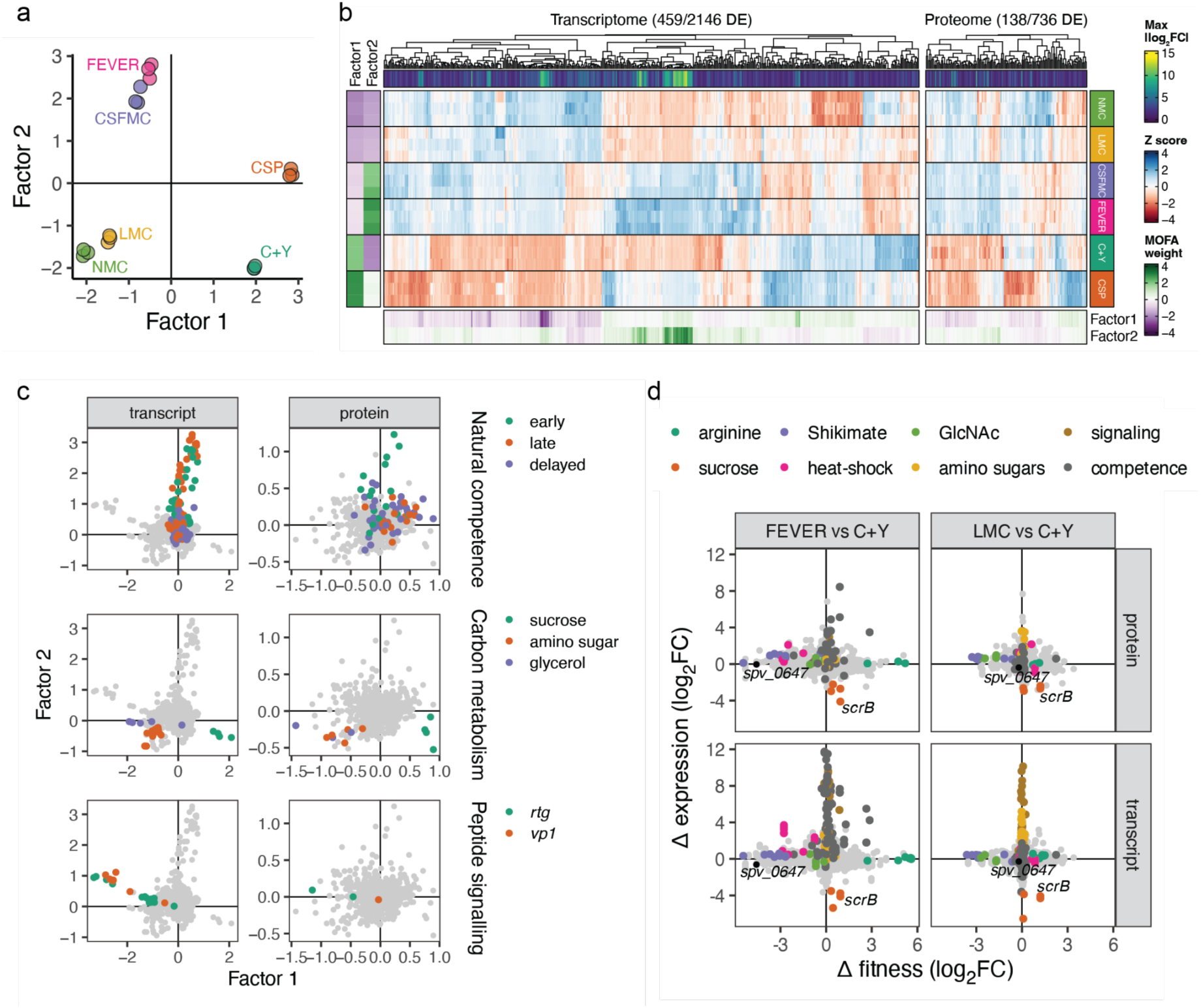
Major differential expression patterns and orthogonality with fitness profiles. (**a**) Dimension reduction by Multi-Omics Factor Analysis (MOFA), considering all, including non-coding, transcripts and the paired nature of the proteome-transcriptome samples. (**b**) MOFA weights correlate well with differential expression patterns between any two growth conditions. (**c**) MOFA uncovers main differential expression programs across all growth conditions. Associations of features with samples can be derived from similarity in weight and direction in this space compared to those of the samples in panel (a). Competence gene classification as defined by Slager and colleagues^20^. (**d**) Orthogonality between differential expression and fitness, illustrated for FEVER and LMC compared to C+Y.

Firstly, natural competence was clearly activated in the CSP, FEVER and CSFMC conditions, distinguishing them from the others (Fig. 5a,c). For these latter two, this can likely be attributed to the relatively high pH in these media (Extended Data Table 1)^57^. The protein response also appeared to be lagged in these two conditions compared to CSP, suggesting slower competence activation (Fig. 5c). Secondly, genes in the *rtg* and *vp1* loci were strongly upregulated in the defined media (Fig. 5c, Extended Data Fig. 4b). These loci both encode a double cell-cell communication loop, where Rgg regulators control glycine-glycine peptide expression^58,59^. Since peptides can also serve as a source of amino acids, we speculate that increases in peptide import might be a response to specific nutrient limitation stimuli. This in turn could kickstart intensive quorum sensing activation as a byproduct. Lastly, genes responsible for the uptake and metabolism of sucrose and amino sugars were indeed associated with the media containing these types of sugars: C+Y / CSP and LMC / NMC, respectively (Fig. 5c, Extended Data Table 1).

Although sucrose import operon *scrAK* was upregulated in the corresponding media, our CRISPRi-seq assay indicated it was not essential for growth, which can be explained by the additional presence of glucose (Extended Data Table 1). In contrast, sucrose catabolism operon *scrBR* was both upregulated and conditionally essential (Fig. 5d). These results imply again toxicity of phosphotransferase sugar import without degradation, as for GlcNAc. Indeed, recent work from our laboratory showed this toxicity can be alleviated by simultaneous knockdown of *scrA* and *scrB*, inhibiting sucrose import and with that likely sugar-phosphate toxicity^35^. These results also imply that, as we showed for GlcNAc, pneumococci import sucrose even in the presence of glucose, at least in sufficient amounts to be toxic in the absence of downstream catabolism.

Strikingly, *nagA*, *nagB* and *manLMN* were not among the many amino sugar metabolism genes upregulated in NMC and LMC. Instead, these mostly comprised genes involved in uptake (*nanP*, *satABC*) and metabolism (*nanA*, *nanB*, *nanE-1*, *nanK*, *nanE-2*) of other amino sugars not present in the media (Fig. 5d). This observation of orthogonality between gene essentiality and expression, whether referring to the transcriptome or proteome, was dominant across the whole genome and all tested conditions (Fig. 5d, Extended Data Fig. 4c). We did find exceptions to this trend, *e.g.*, the example of *scrB* given above, or the conditional upregulation and essentiality of the HrcA-regulated heat-shock protein-encoding operon at 40°C, the latter of which was not observed on the proteome level presumably due to the short time between heat exposure and sample processing (5 min) (Fig. 5d). In general, however, our results indicated that differentially expressed genes were rarely differentially essential and *vice versa* (Extended Data Fig. 4c), which fits observations made by others in the same and other organisms^12–17^.

## Discussion

In this work, we investigated how pneumococci adapt to different environmental settings in terms of genome-wide gene fitness effects and expression, both on the transcript and protein level. This allowed us to not just re-assess known relationships between these regulatory layers in *S. pneumoniae*, but also to work out specific molecular responses to concrete stimuli. Global patterns roughly corroborated biology as it is known in other organisms: transcriptomes correlate moderately with proteomes^8–11^, and either appear almost entirely statistically independent from genome-wide fitness impact^12–17^. However, Jensen and colleagues noted this does not necessarily imply biological independence: genes relating to the same general pathways could be either differentially essential, or differentially expressed, and as such still coordinated^12^. We see one such example, where *nagA*, *nagB* and *manLMN* are required for growth on GlcNAc, but not differentially regulated, whereas other amino sugar metabolism genes are, while they are not essential. The data suggest that basal expression levels mostly suffice for bacterial survival in distinct environments, rendering differential expression of essential genes generally redundant. Conversely, differential expression does not signal a changed need for a functional gene copy *per se*: for instance, a gene can be upregulated in one growth condition compared to another, but essential in both. Moreover, expression-fitness correlations could be masked by genetic redundancy, which could be addressed by multiplexed knockdown approaches such as dual CRISPRi-seq^35,60^. In addition, it is important to acknowledge that the selective pressures that shaped regulation of certain genes in response to specific cues in the natural habitat might be absent in our artificial laboratory conditions, whereas those cues can still be present, potentially leading to differential expression without a difference in essentiality. Our data indicate that expression and fitness effects provide almost completely mutually complementary information on how bacteria adapt to different environments. This in turn makes the case for multi-omics approaches to understand how bacteria deal with their surroundings and stresses, including for instance antibiotic pressures, with potentially important implications for treatment of infections. It also suggests that differential expression assays by themselves are not necessarily the best tool to uncover potential therapeutic targets, as they might not point to essentiality.

An environmental factor that is inherently intertwined with infection and disease, is ambient temperature. As *S. pneumoniae* moves from the nasopharynx (32°C) to other parts of the body (37°C), where it can bring about fever (>38°C), it faces considerable temperature shifts^61^. This is known to affect the cells in a myriad of ways, including membrane properties such as fluidity and permeability. Bacteria are known to counter these effects through homeoviscous adaptation, *i.e.*, by adjusting the relative levels of different fatty acid types in the membrane^45,46^. Previous studies indicated that such adaptation by *S. pneumoniae* in response to temperature changes was independent of fatty acid biosynthesis master regulator FabT^37^. Here, we report that *spv_0647* encodes a transcriptional regulator that enables pneumococci to maintain proper saturated:unsaturated fatty acid (SFA:UFA) balance, critical during heat stress, by means of mediating *fabM* and potentially *fakB3* transcription. As such, we named this gene *fasR*, for fatty acid saturation regulator. Since we did not find putative binding sites for FasR in the promoter regions of differentially expressed genes using motif enrichment analyses with the bioinformatic MEME suite^62^, future research may focus on establishing DNA binding sites and sequences using, for example, EMSA and ChIP-seq. In addition, it could be insightful to examine lipid head groups, which might also influence membrane properties but are missed by GC-FAME, the lipid analysis technique used here. Generally, metabolomics techniques could provide an additional layer of information, as it would be possible to gauge for instance the abundance of lipid intermediates, further narrowing down the enzymes potentially regulated by FasR. Moreover, most of the observed differences between the growth conditions tested here were metabolic in nature, and could potentially be better understood in the light of metabolomic profiles.

We elaborated on one such example here, showing that N-acetylglucosamine (GlcNAc) degradation after uptake is essential for pneumococci, and corroborating this is the case for sucrose as well. As both sugars are phosphorylated upon import, we hypothesize these are instances of sugar-phosphate stress^30^. Moreover, these toxicities are seen in the presence of glucose, suggesting simultaneous uptake of these sugars. Indeed, we showed a slowdown in glucose uptake in the presence of GlcNAc while the growth rate of WT cells was unaffected. Although this challenges the dogma of CcpA-based glucose preference in *S. pneumoniae*^26,27^, it may not be illogical from an evolutionary perspective, as glycans such as GlcNAc are far more available for scavenging in its natural niche, the human nasopharynx, than glucose is^63–65^.

Although we provide a broad overview of the large, systems level data compendium presented here, we have only worked out a few processes that stood out in detail. Many more biological insights may be concealed in these data and we encourage the community to use them to their advantage. To facilitate such efforts, we have also integrated our genome-wide fitness, transcript and protein data sets with our recently renewed genome browser PneumoBrowse 2 (https://veeninglab.com/pneumobrowse)^23^. On top of that, we highlighted some of these avenues to be explored, such as the activation of quorum sensing systems in nutrient-limiting conditions, or the potential fitness gain upon arginine retention during heat stress. Despite the fact that heat itself has been shown to influence CRISPRi efficiency^66^, we clearly do retrieve the standard core essentialome in our high-temperature growth condition and are therefore confident regarding data quality.

Our findings on GlcNAc metabolism and membrane homeostasis contribute towards our knowledge of gene function and the biology behind environmental adaptation. We note that many pneumococcal genes remain of unknown function, and that adaptation is a complex phenotype, brought about on multiple, interacting regulatory levels. *S. pneumoniae* is notoriously versatile in terms of niche adaptation, as it can occupy many micro-environments in the human body^1–3^. A deeper understanding of the biology underpinning this capacity, in the pneumococcus as well as other microorganisms, therefore ultimately also yields a deeper understanding of human health and disease.

## Methods

### Bacterial strains and growth conditions

*Streptococcus pneumoniae* D39V serotype 2 and derivatives were routinely grown at 37°C in C+Y liquid medium (pH 6.8) without shaking, or on Columbia agar plates with 2.5-5% (v/v) defibrinated sheep blood (CBA, Thermo Scientific) at 5% CO2, supplemented with 0.5 μg·mL^-1^ erythromycin, 0.5 μg·mL^-1^ tetracycline or 50 ng·mL^-1^ anhydrotetracycline when appropriate. Strains used in this study are listed in Supplementary Table 18. Other growth conditions and media were prepared as described by Aprianto and colleagues (2018)^7^, with the only adaptation of continuous culturing at 40°C in the FEVER condition for the CRISPRi-seq assay. Growth assays for sugar preference were performed in a C+Y liquid or agar plate background without added sugars, supplemented with 9.4 mM glucose, 9.4 or 0.94 mM N-acetyl-D-glucosamine (Sigma-Aldrich, A3286), or both sugars as appropriate.

### Mutant strain construction

Donor DNA constructs carrying the insert of interest with ∼1000 bp flanking regions homologous to the insertion site were produced with a one-pot Golden Gate assembly strategy using Type II restriction enzymes BsaI, Esp3I or SapI (New England Biolabs)^67^. Restriction sites were introduced via the primers during PCR amplification. Used oligonucleotides and restriction enzymes are listed in Supplementary Table S19. We used the PT5-3 variant of the Ptet promoter as characterized by Sorg and colleagues (2020)^68^.

Subsequent transformation with the donor DNA was performed as described previously^18^.

Briefly, pneumococci were cultured at 37°C to early exponential phase (OD595 ∼0.1) followed by addition of 0.1 μg·mL competence-stimulating peptide 1 (CSP-1) and growth for another 12 min to activate competence. 100 μL activated culture was mixed with donor DNA at a concentration of 1 ng·μL and further cultured at 30°C for 20 min. 900 μL fresh C+Y was added, and the culture was cultivated at 37°C for another 1.5 h for the transformants to recover. The culture was plated and incubated overnight as described above. Colonies were re-streaked on plates and incubated overnight again. Transformant colonies were picked and grown in liquid C+Y until OD595 ∼0.3, and stocked in 14-20% glycerol. Genotypes of mutant strains were confirmed by Sanger sequencing (Microsynth).

### Growth curves

Pre-cultures were grown in C+Y to OD595 ∼0.1 at 37°C, diluted 100× in the appropriate medium and loaded into flat-bottom 96-wells plates at 250 μL per well. 50 ng·mL-1 anhydrotetracycline was added to both pre-cultures and dilutions where appropriate. Experiments were always performed in triplicate (three separate pre-cultures), unless specifically stated in the figure. For each replicate, a representative curve was chosen out of three technical replicates (wells within the same plate), on the basis of most frequent appearance between the other two technical replicates across all time points. Blanks were added to each plate to assess potential contamination. In experiments where glucose levels were also measured, the number of technical replicates equaled the number of glucose measurement time points to allow subsampling for that purpose. Optical density was measured every 10 minutes at 595 nm in a plate reader (Tecan MPlex, F200 or M200 series). In temperature variation experiments, three identical plates were prepared and measured simultaneously in three different plate readers, each set at a different temperature (30°C, 37°C or 40°C). Plates were sealed with parafilm in this case to avoid excessive evaporation. Raw OD values were normalized per well to the theoretical start OD of 0.001 by subtraction, and lower values were also set to this theoretical minimum.

### Glucose oxidation assays

Samples were obtained by pausing the plate reader during growth curves assays at time points indicated in the figures, transferring the contents of technical replicates (200 μL) to microtubes, and resuming OD measurements as fast as possible. Microtubes were immediately spun down on a tabletop mini centrifuge for 3 min to pellet the cells, after which 150 μL supernatant was transferred to a new microtube. The first time point sample (0 h) was taken directly from the growth curve pre-culture and treated in the same way. Samples were snap frozen using liquid nitrogen and stored at -80C until glucose measurements.

Glucose concentrations were measured with a Glucose (GO) Assay Kit (Sigma-Aldrich GAGO20) according to instructions of the manufacturer, except for the total sample volumes. We scaled all volumes down so that samples were 150 μL instead of 5 mL, allowing for higher-throughput measurements using flat-bottom 96-wells plates in a Tecan plate reader, instead of cuvettes.

### Proteo-transcriptomics

Wild-type cells were pre-cultured in each respective growth medium to OD600 ∼0.1 and diluted to OD600 ∼0.05. CSP cultures were supplemented with 0.1 μg·mL-1 CSP-1 for 20 min and FEVER cultures were transferred to 40°C for 5 min. Each of three replicates was split into two subsamples, one of which was subjected to RNA-seq and the other to quantitative LC-MS.

Total RNA was extracted and cDNA libraries were constructed as before, without rRNA depletion^7^. Libraries were sequenced on an Illumina NextSeq machine at GeneCore, EMBL Heidelberg. Read quality was checked with FastQC (v0.11.5, https://www.bioinformatics.babraham.ac.uk/projects/fastqc/) before and after trimming off

TruSeq3 adapters, leading and trailing bases below a phred score of 3, cutting regions if average phred scores went below 20 in a sliding window of 5 bases and only keeping reads with a minimum length of 50 bases using Trimmomatic (v0.36)^69^. Reads were aligned to the *S. pneumoniae* D39V reference genome (CP027540) using STAR (v2.5.3a)^70^ and reverse strand transcripts were counted with featureCounts (Subread v1.5.3)^71^, using multi-mapping, overlap and fractional count modes, as before^7^. Downstream analyses were done with DESeq2 (v1.34.0)^34^ in R (v4.1.1), where differential expression was tested against an absolute log2 fold change of 1 at an alpha of 0.05, and counts were normalized for Principal Component Analysis and Multi-Omics Factor Analysis with a blind rlog transformation^34^. Transcripts per million were calculated per sample with *n* genes for each gene *i* as: 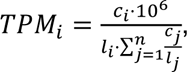, where *c* and *l* are vectors of length *n* representing the raw transcript counts and lengths, respectively. Cells in the proteomics subsamples were lysed with a bead beater and further treated for LC-MS at the Proteomics - Mass Spectrometry Service Facility, University of Groningen. Briefly, sample volumes were reduced by freeze drying, protein concentrations were determined with a BCA assay (Thermo, 23252) and samples were reconstituted in a 100 mM ammonium bicarbonate buffer. Alkylation of 100 μg protein was achieved by adding iodoacetamide to a final concentration of 40 mM and incubation for 45 min at room temperature, in the dark. Samples were diluted 2× in 100 mM ammonium bicarbonate and overnight digestion was performed at 37°C, 400 rpm with mass spectrometry grade trypsin (Promega, V5280) using a 1:50 trypsin:protein (μg:μg) ratio. The reaction was stopped by adding trifluoroacetic acid to a final concentration of 1%. Pierce® C18 tips (Thermo, 87784) were used for sample cleanup by solid phase extraction according to the manufacturer’s instructions. The elute fraction was dried under vacuum and reconstituted with 20 μL 2% acetonitrile, 0.1% formic acid (FA). Peptide separation was performed with 2 μL peptide sample using a nano-flow chromatography system (Thermo, EASY nLC II) equipped with a reversed phase HPLC column (75 μm, 15 cm) packed in-house with C18 resin (Dr. Maisch, ReproSil-Pur C18–AQ, 3 μm resin) using a linear gradient from 95% solvent A (0.1% FA, 2% acetonitrile) and 5% solvent B (99.9% acetonitrile, 0.1% FA) to 28% solvent B over 90 min at a flow rate of 200 nL·min^-1^. The total MS time was 120 min. The peptide and peptide fragment masses were determined by an electrospray ionization mass spectrometer (Thermo, LTQ-Orbi-trap XL). Peptides were mapped to the *S. pneumoniae* D39V (CP027540) protein fasta file and quantified as both label-free quantification (LFQ) and intensity Based Absolute Quantification (iBAQ) values with MaxQuant^72^, using a false discovery rate (FDR) cutoff of 0.01. Downstream analyses were performed with R package DEP (v1.16.0)^73^ using LFQ values as input, where a protein was considered differentially enriched if its absolute log2 fold change was significantly greater than 1, with an FDR-adjusted P-value below 0.05. We retained only proteins that were detected in at least two out of three replicates per condition, normalized with a variance-stabilizing transformation, and imputed missing values with the MinProb method as implemented in DEP, since values were not missing at random, but biased towards lower intensities (Extended Data Fig. 3). Gene Ontology (GO) term enrichment analysis was carried out using R package clusterProfiler (4.2.2)^74^. For visualization purposes we normalized for library size bias per sample as proteins per million: , akin to TPM as described above.

We used the blind rlog and variance-stabilized quantifications of transcripts and proteins with non-zero variance across samples as input for Multi-Omics Factor Analysis using the R package MOFA2 (v1.4.0)^56^ with default settings.

### RNA-seq Δ*spv_0647*

Wild-type (VL1) and Δ*spv_0647* (VL6297) strains (Supplementary Table 18) were pre-cultured in C+Y medium at 30°C and 37°C until OD595 ∼0.3. Pre-cultures were diluted to OD595 ∼0.01 in quadruplicates and grown at 30°C and 37°C until OD595 ∼0.3-0.4. Cells were pelleted by centrifugation (4°C, 10000 *g*, 5 min), supernatants removed, and pellets were stored at -80°C after snap-freezing with liquid nitrogen.

Total RNA was isolated with a High Pure RNA Isolation Kit (Roche, 11828665001) as before with minor adaptations^7^. Briefly, the pellets were resuspended in 400 μL Tris-EDTA buffer and transferred to tubes with 50 μL SDS 10%, 500 μL phenol-CHCl3 and glass beads. Cells were lysed with a bead beater (3× 45 s with 45 s breaks) and pelleted by centrifugation (4°C, 21000 g, 15 min). 300 μL of the aqueous phase was mixed into 400 μL lysis/binding buffer. The samples were loaded onto columns and centrifuged (8000 g, 30 s). 100 μL DNase mix (90 μL buffer, 10 μL DNase I) was loaded onto the column filter and incubated for 1 h at room temperature. Samples were washed once with wash buffer I (500 μL) and twice with wash buffer II (first time 500 μL, second time 200 μL) by centrifugation (8000 g, 30 s). Samples were eluted in 50 μL elution buffer and incubated for 10 min at room temperature. Sample quality was checked by NanoDrop and Fraction Analyzer (Agilent Technologies), and samples were stored at -80°C. Samples had RNA Quality Numbers (RQN) between 5.7 and 7.8.

Per strain-temperature combination, the three samples with minimal potential gDNA contamination and the highest RQN were selected for cDNA library preparation and sequencing at the Genomic Technologies Facility, University of Lausanne. Briefly, RNA-seq libraries were prepared from 100 ng of total RNA with the Illumina Stranded mRNA Prep reagents (Illumina) using a unique dual indexing strategy and following the official protocols. The polyA selection step was replaced by an rRNA depletion step with RiboCop for Bacteria, mixed bacterial samples, reagents (Lexogen). Libraries were quantified by a fluorometric method (QubIT, Life Technologies) and their quality was assessed on a Fragment Analyzer (Agilent Technologies). Sequencing was performed on an Illumina NovaSeq 6000 for 100 cycles, single read. Sequencing data were demultiplexed using the bcl2fastq2 Conversion Software (v2.20, Illumina).

Read quality was checked with FastQC (v0.11.9) and MultiQC (v1.15)^75^ before and after trimming off leading and trailing bases below a phred score of 3, cutting regions if average phred scores went below 20 in a sliding window of 5 bases and only keeping reads with a minimum length of 50 bases using Trimmomatic (v0.36)^69^. Reads were aligned to the *S. pneumoniae* D39V reference genome (CP027540) using bowtie2 (v2.4.5)^76^, using soft-clipping (with the “--local” option) to account for any remaining partial adapters.

Transcript counts were extracted with featureCounts (v2.0.6)^71^ and downstream analyses were performed with DEseq2^34^ in R, as described for the proteo-transcriptome experiment.

### CRISPRi-seq

An IPTG-inducible genome-wide CRISPRi library was used as described before^18,19^. Briefly, the library was pre-cultured in the respective growth conditions (Extended Data Table 1) to OD595 ∼0.1, followed by 100× dilution supplemented with 1 mM IPTG for CRISPRi induction and CSP-1 (0.1 μg·mL^-1^), when appropriate, to yield quadruplicates for each growth condition with and without IPTG. Samples were grown to OD595 ∼0.1 once (LMC, NMC, CSFMC, FEVER) or twice (C+Y, CSP, THY, BMC) by 100× back-dilution, corresponding to 7-14 generations of exponential, sample-wide growth. Pellets were harvested on ice, centrifuging twice (4°C, 15 min at 4000 *g* and 5 min at 12000 *g*, respectively), discarding supernatant and resuspending in PBS. gDNA isolation, library preparation and sequencing were done as described in our published protocols^18^. CRISPRi-induced NMC samples 54 and 55 got mixed up during gDNA isolation and were henceforth regarded as technical replicates.

Samples were split over two sequencing runs on an Illumina MiniSeq system with our published custom sequencing protocol^18^. sgRNA counts were extracted with 2FAST2Q (v2.5.2)^77^ using default settings (minimal phred score 30, one mismatch allowed). Differential fitness analyses were performed with DESeq2 (v1.34.0)^34^ in R (v4.1.1), testing against a minimal (difference in) absolute log2 fold change of 1 at an alpha of 0.05 for statistical significance. We collapsed the induced NMC technical replicates into one biological replicate with the DESeq2 function collapseReplicates().

Given growth at a rate of 2^n^, fitness quantifications on a log2FC should scale linearly across conditions. We observed this was not the case between conditions grown for a different sample-wide number of generations, implying a strong library composition bias and rendering DESeq2-based interaction effects uninformative. We corrected this bias by estimating generation numbers per CRISPRi strain in each induced sample, linearizing them over all samples with a LOESS transformation, and recomputing corresponding sgRNA counts. These were rounded and used as DESeq2 input. Sample-wide CRISPRi induction generation numbers (0, 7 or 14) were scaled and centered with the built-in R function scale(), and together with growth condition and their interaction term used as explanatory variables in the DESeq2 design formula. So, default DESeq2 methods were applied after we normalized for the non-linear generation effect.

Specifically, sample-wise raw sgRNA counts *c* were first normalized for library size bias: 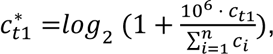, where *t1* indicates counts after growth (as measured). We used uninduced sample counts as a proxy for the relative starting distribution of CRISPRi strains per sample prior to induction, i.e., at *t0*. Given bacterial growth occurs at a rate of *2^m^*, so that in general 𝑐_,$_ = 𝑐_,%_ ⋅ 2^0^, the relative counts per strain at *t0* are: 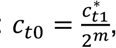, where in our case *m* is 7 or 14. To obtain a more robust estimate, we averaged *ct0* per strain across each set of four uninduced replicate samples, yielding 𝑐,_%_. This represents an estimate of the relative starting counts per CRISPRi strain, or sgRNA, for all samples within a given growth condition. Following the same standard growth equation as above, the relative counts after growth should then equal 𝑐^∗^ = 𝑐,_%_ ⋅ 2^1^, and so we estimated the strain-wise generation numbers in each induced sample as: 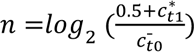, where a pseudocount of *0.5* was added to avoid zeroes. Together, this yields the generation number matrix **N**, where rows represent CRISPRi strains, and columns induced samples of all growth conditions. Generation number estimates were then linearized with the cyclicloess method of the normalizeBetweenArrays() function from the limma R package (v3.50.1)^78^, with **N** as input and otherwise default parameters. The resulting matrix columns represent the corrected generation number estimates *n\** per induced sample, which we then used to estimate relative strain counts after growth per induced sample following the same standard growth dynamics described above: 𝑐2 = 𝑐2 ⋅ 2^1∗^, where Lastly, we scaled these corrected counts to their original library size as: 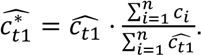 These normalized sgRNA counts for the induced samples were subsequently used as DESeq2 input for downstream analyses, together with the original sgRNA counts of the uninduced samples.

### Foldseek

Foldseek^47^ was accessed through the online submission portal (https://search.foldseek.com) and ran with UniProt accession number A0A0H2ZQ31, for *spv_0647*. Only matches with crystalized protein structures from the PDB100 database were considered for this work.

### Dual CRISPRi-seq analysis

Fitness scores (log2FC values) for every unique pair of 869 sgRNAs were obtained from Dénéréaz and colleagues^35^. Scores of sgRNAs in combination with themselves (twice the same sgRNA in the same CRISPRi strain) were used as baseline fitness estimates of the corresponding sgRNA, as in the original study.

### GC-FAME

Strains were pre-cultured at 30°C, 37°C or 40°C with or without 50 ng·mL^-1^ anhydrotetracycline and 1 mM IPTG, as appropriate, to an OD595 ∼0.3 and concentrated 10× by centrifugation (3 min, 8000 *g*). Quadruplicate samples were grown from these pre-cultures in the same growth conditions, as appropriate, in volumes of 45 mL per sample to OD595 ∼0.2-0.3. Samples were kept at their respective growth temperatures throughout the experiment. Cells were pelleted by centrifugation (15 min, 1968 *g*), resuspended in ∼1 mL of the remaining medium after discarding supernatant, transferred to 2 mL screw-cap tubes and centrifuged again (5 min, 20238 *g*). Cells were washed by supernatant removal, resuspension in 1 mL PBS and centrifugation (5 min, 20238 *g*). Supernatant was removed again, and pellets were stored at - 80°C following snap-freezing with liquid nitrogen.

For derivatization of fatty acid methyl esters, frozen cell pellets were resuspended and then vortexed in 100 µL concentrated sulfuric acid (96%) and 200 µL methanol. Samples were then boiled by incubation in boiling water for 5 min and allowed to cool to RT. 300 µL of dichloromethane was added, samples were then vortexed and centrifuged 1 min at 15000 *g*.

The organic (bottom) layer was transferred to a new tube containing a pinch of Na2SO4 (to remove any remaining water), mixed by vortexing, and centrifuged 1 min at 15000 *g*. The supernatant was then transferred to an HPLC tube and stored at 4°C until being loaded on the GC-MS. 1 µl of sample was injected into an Agilent 7890 Gas Chromatograph equipped with an Agilent G3903-63011 column, an Agilent 5977A Mass Detector and an Agilent 7693 Autoinjector. The carrier gas was helium, and the column oven temperature program was the following: 150°C for 0.5 min; ramp temperature 25°C/min to 230°C then hold for 1 min; ramp temperature 5°C/min to 245°C then hold for 1 min. Spectra from 50-500 m/z were collected after a 4 minute solvent delay. The ion source temperature was 230°C.

Raw data was exported as CDF files, which were filtered to remove empty scans and converted to .mzML files using MZmine 4 (mzio.io)^79^. Spectral alignment, deconvolution, and relative abundance analyses were performed using MS-Hub as part of the Global Natural Products Social Molecular Networking (GNPS) GC-MS EI Data Analysis pipeline^80^. Peaks representing methylated fatty acids were identified based on comparison of spectra and retention times to those obtained using TraceCERT 37 component fatty acid methyl ester standard (Sigma Aldrich) across the same instrumentation protocol. For each sample, peak intensities were summed per unique fatty acid. These values were standardized to sum to 100 per sample, and analyzed with the R package compositions (v2.0.8) to account for the particular biases and inherent complexities of compositional data^55^. Specifically, we used the acomp() function to transform the data for Principal Component Analysis, and the irl transformation in combination with a compositional (mlm) ANOVA for hypothesis testing.

## Data availability

For the proteo-transcriptomic profiling, raw RNA-seq data is available on SRA, accession number PRJNA527271, and LC-MS data on the UCSD MASSive repository, accession number MSV000097932. Raw CRISPRi-seq data is available on SRA, accession number PRJNA1262882. RNA-seq data for the mutant experiment can be found on SRA, accession number PRJNA1262992, and GC-FAME data on the UCSD MASSive repository, accession number MSV000097619.

### Acknowledgments

We would like to thank all members of the Veening lab for fruitful discussions, Andrew Quinn for advice on the use of glucose oxidation assays, Florian Bock for practical biochemistry guidance, Johann Mignolet and Julien Dénéréaz for providing genetic constructs, Axel Janssen for integrating our data with PneumoBrowse 2, Kathrin Fröhlich for bringing the phenomenon of sugar-phosphate stress to our attention, and James Saenz for valuable insights regarding membrane properties. Moreover, we thank Rieza Aprianto for proteo-transcriptomic sample preparation, and Johan Hekelaar at the Proteomics Facility of the University of Groningen for the LC-MS quantifications and his help in proteomics analyses. RNA-seq for the proteo-transcriptome experiment was performed at GeneCore, EMBL, Heidelberg. Library preparation and RNA-seq of the spv_0647 mutant and WT strain were performed at the Lausanne Genomic Technologies Facility, University of Lausanne, Switzerland. This work was supported by Swiss National Science Foundation (SNSF) PostDoc Mobility fellowship P500PB_225439, ERC consolidator grant 771534, SNSF grants 310030_192517, 310030_200792 and NCCR ‘AntiResist’ 51NF40_180541, and NIH/NIDCR R00-029228.

## Author contributions

VdB performed experiments, analyzed data, and wrote the manuscript. XL supervised experiments. JT and MB performed experiments and analyzed data. JLB supervised experiments, analyzed data and edited the manuscript. JWV supervised the study and edited the manuscript.

## Materials & Correspondence

All correspondence should be directed to JWV.

## Extended data

**Extended Data Table 1.** Growth conditions used in this study as defined in Aprianto *et al.* (2018).

|  | Niche-Mimicking Condition | Temp. (°C) | pH <sup>a</sup> | Sugar | Experiment <sup>b</sup> | CRISPRi generations <sup>c</sup> |
| --- | --- | --- | --- | --- | --- | --- |
| NMC | Nasopharynx | 30 | 7 | GlcNAc | T, P, F | 7 |
| LMC | Lungs | 37 | 7 | GlcNAc | T, P, F | 7 |
| CSFMC | Cerebrospinal fluid | 37 | 7.8 | Glc | T, P, F | 7 |
| FEVER | Meningitis fever | 40 | 7.8 | Glc | T, P, F | 7 |
| C+Y |  | 37 | 6.8 | Glc + Scr | T, P, F | 14 |
| CSP |  | 37 | 6.8 | Glc + Scr | T, P, F | 14 |
| BMC | Blood | 37 | 7.4 | Glc | F | 14 |
| THY |  | 37 | 7.5 | Glc | F | 14 |

<sup>a</sup> pH was calibrated at the start of the experiment.
<sup>b</sup> Column indicates which omics layers were measured.
<sup>c</sup> Column indicates for how many overall generations the CRISPRi pool was grown as estimated from the whole culture OD<sub>595</sub>.
GlcNAc: N-acetylglucosamine, Glc: glucose, Scr: sucrose, T: transcriptome, P: proteome, F: fitnessome.

**Extended Data Fig. 1.**
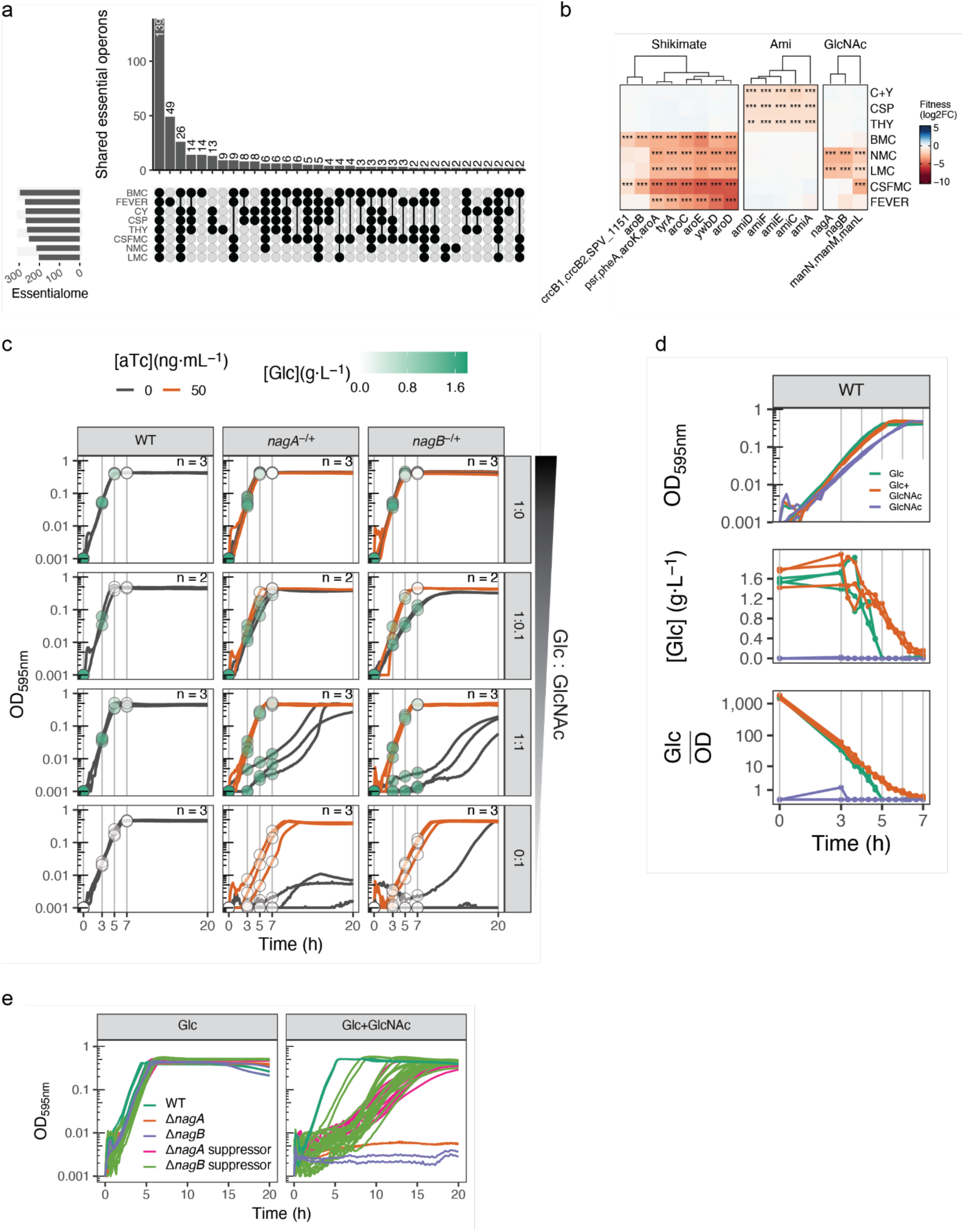
CRISPRi-seq profiles with focus on metabolism. (**a**) General overlap of essentialomes between growth conditions: number of sgRNA targets classified as essential (log2FC<-1, Padj<0.05). (**b**) Selected fitness effects of sgRNAs targeting members of the Shikimate pathway, Ami peptide transport system and N-acetylglucosamine (GlcNAc) import and catabolism genes across growth conditions. (**c**) Glucose concentrations in the medium during growth of WT strains and aTc-inducible ectopic complementation Ptet-*nagA* and Ptet-*nagB* strains in which the native genes are deleted, respectively (*nagA*^-/+^, *nagB*^-/+^). Glucose quantifications are superimposed on the corresponding sample growth curves for which they were measured. Media contained varying glucose:GlcNAc molar ratios, as indicated. Biological replicates are displayed in the relevant panels. (**d**) Glucose concentrations in the medium, measured every 20 min, during growth of WT cells in glucose, GlcNAc, or an equimolar mix of the two. Concentrations and OD were measured on each of three biological replicates and their fraction is shown in the bottom panel. (**e**) Growth curves of WT, *nagA* and *nagB* deletion mutants and all the suppressor isolates of the mutants, grown on glucose or an equimolar mix of glucose and GlcNAc.

**Extended Data Fig. 2.**
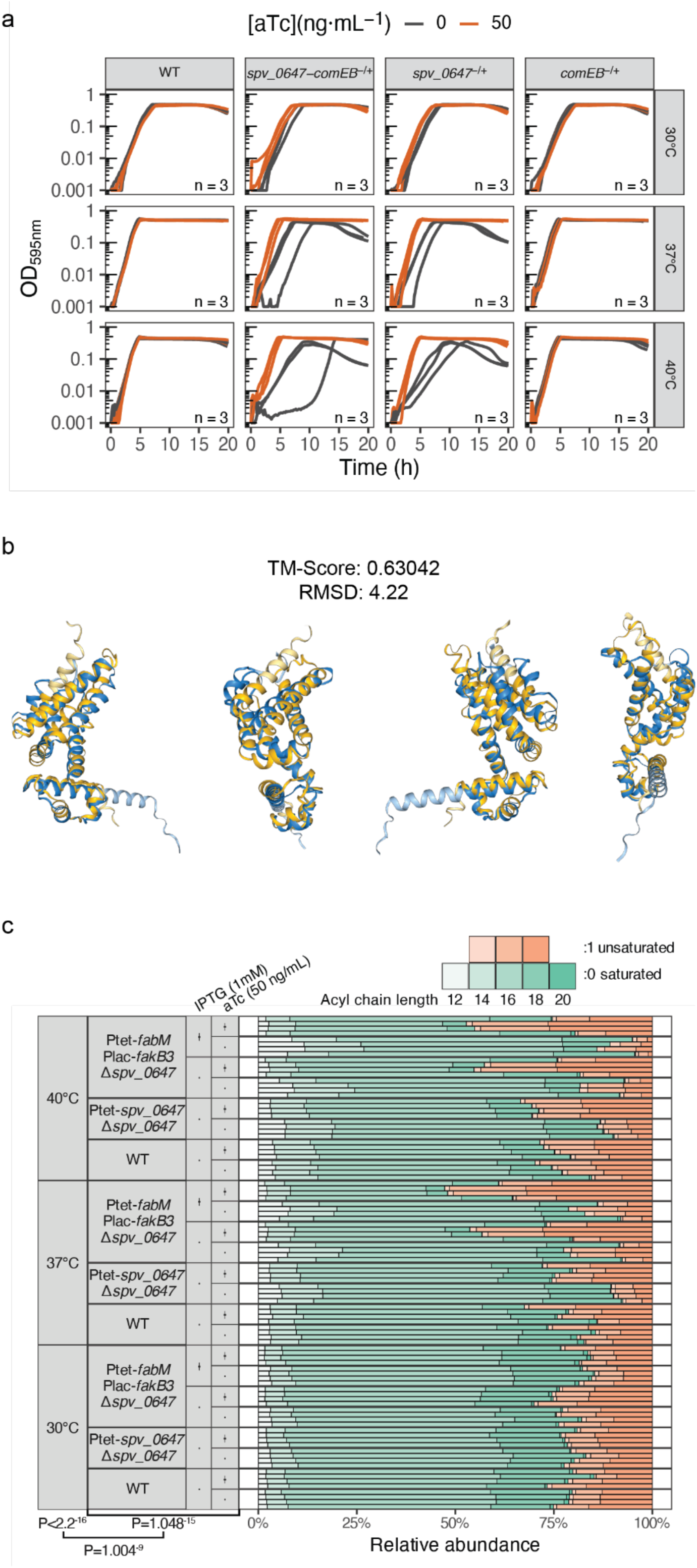
spv_0647 confers a fitness advantage during heat stress and influences fatty acid composition of the membrane. (**a**) Growth curves of the WT strain, and *spv_0647*, *comEB* and the whole operon deletion mutants with aTc-inducible, ectopic complementation, grown at three different temperatures. Biological triplicates are shown. (**b**) Structural alignment of the FoldSeek top hit PDB ID 4MK6 (yellow) and SPV_0647 AlphaFold prediction (blue). (**c**) Full fatty acid membrane composition profiles of all temperature - strain combinations tested. P-values are derived from a compositional (mlm) ANOVA, performed using the R package compositions, based on an irl-transformation^55^.

**Extended Data Fig. 3.**
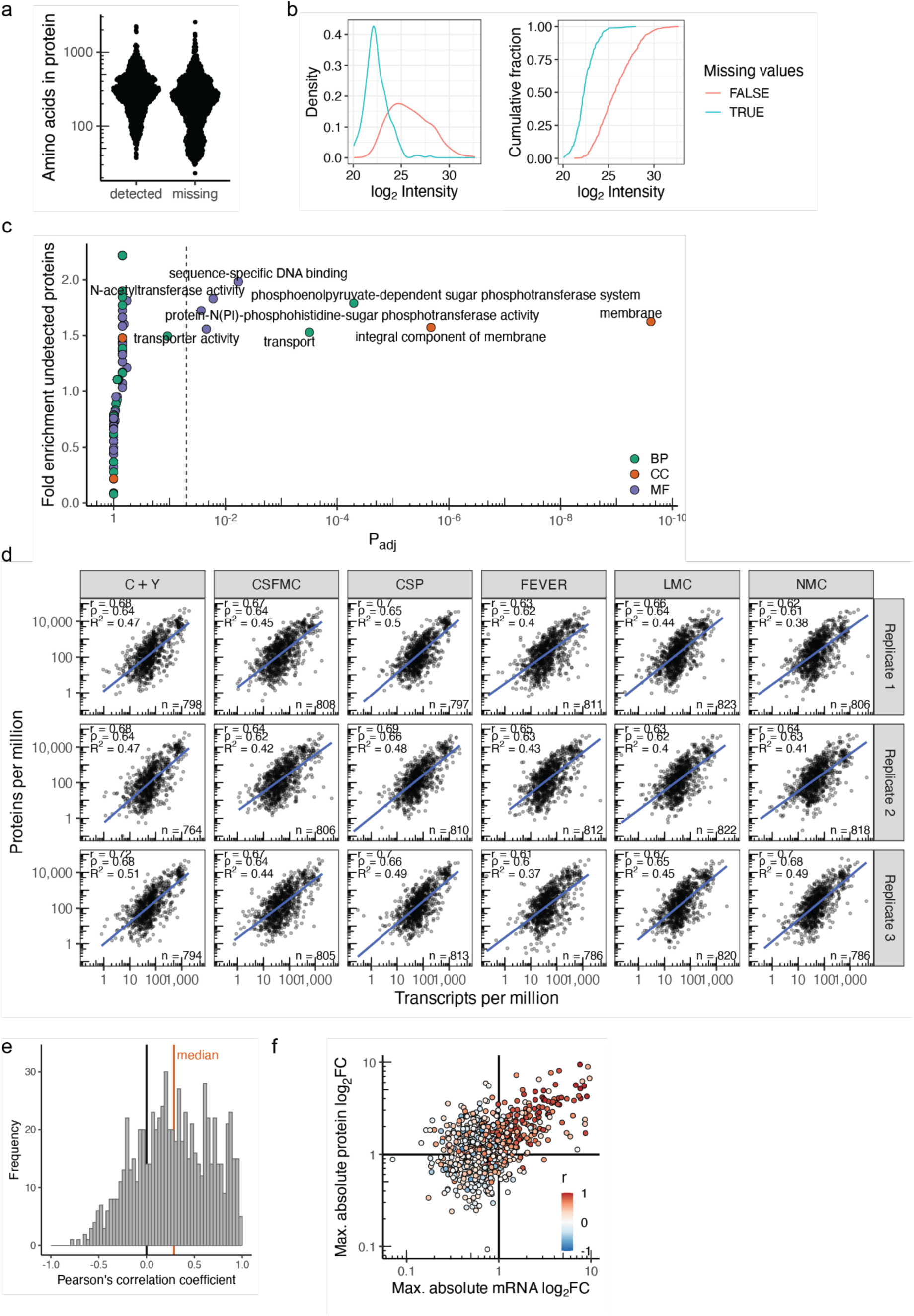
Transcriptome-proteome relationships across growth conditions. (**a**) Shorter proteins were underrepresented in the data set. (**b**) Lowly abundant were detected less frequently, presumably largely due to being close to the detection limit^73^. (**c**) GO term enrichment analysis shows membrane proteins were overrepresented in the set of proteins left undetected. Fold enrichment is the gene ratio (fraction of undetected proteins in the GO set divided by the total of undetected proteins) divided by the background ratio (fraction of total proteins in the GO set divided by the total number of proteins). (**d**) Correlations between relative transcript and protein numbers per sample. The number of genes for which both levels were reliably quantified is displayed in each panel. (**e**) Distribution of correlation coefficients between transcript and protein levels across samples, per gene. Transcript numbers were normalized with the rlog transformation^34^, and protein levels by vsn transformation following imputation of LFQ intensities^73^. (**f**) Maximum differential expression for each between any two growth conditions on the transcript and protein level. Correlations between transcript and protein levels across conditions as in panel (e).

**Extended Data Fig 4.**
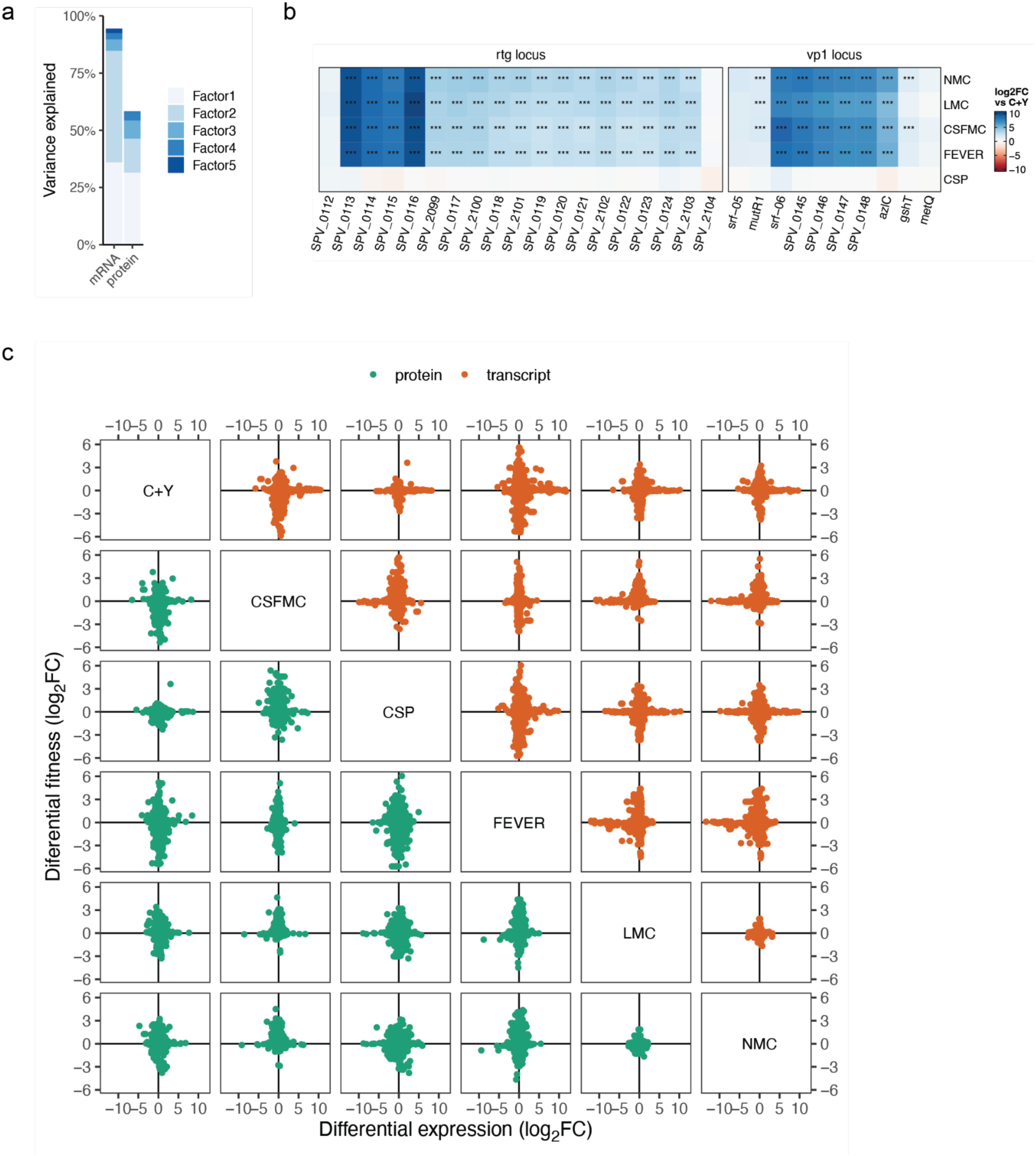
Integrated proteo-transcriptomic profiles. (**a**) Variance explained per dimension in either dataset by Multi-Omics Factor Analysis (MOFA)^56^. (**b**) Differential expression of selected transcripts with high MOFA weights, compared to C+Y. Three asterisks indicates Padj<0.001 for |Δlog2FC|>1. (**c**) Comparison of differential expression (upper triangle: transcriptome, lower triangle: proteome) and fitness quantifications between each pair of growth conditions. Genes targeted by the same sgRNA (i.e., those sharing operons) were assigned the same fitness score.

