## Supplementary Tables 18-19 for "Multi-omics profiling reveals atypical sugar utilization and identifies a key membrane composition regulator in *Streptococcus pneumoniae*"

### Multi-omics profiling reveals atypical sugar utilization and identifies a key membrane composition regulator in *Streptococcus pneumoniae*

Vincent de Bakker, Xue Liu, Jonah Tang, Matthew Barbisan, Jonathon L. Baker, Jan-Willem Veening

**Supplementary Table 18. Bacterial strains that were used in this work.**

| ID | Genotype | Source |
|---|---|---|
| VL1 | Wild type *Streptococcus pneumoniae* D39V serotype 2 | Slager *et al.* (2018) [1] |
| VL1998 / DCI23 | D39V, bgaA::Plac-dcas9sp (tet), prs1::PF6-lacI (gen) | Liu *et al.* (2017) [2] |
| VL2210 | D39V, prs1::PF6-tetR (gen) | Liu *et al.* (2021) [3] |
| VL4012 | D39V, bgaA::Ptet (tet) | Lab collection |
| VL6009 | D39V, prs1::PF6-tetR (gen), bgaA::Ptet-nagA (tet), ΔnagA (ery) | This study |
| VL5893 | D39V, prs1::PF6-tetR (gen), bgaA::Ptet-nagB (tet), ΔnagB (ery) | This study |
| VL5305 | D39V, ΔnagA (ery) | This study |
| VL5306 | D39V, ΔnagB (ery) | This study |
| VL7271 | ΔmanLMN (kan) | This study |
| VL7272 | ΔnagA (ery), ΔmanLMN (kan) | This study |
| VL7273 | ΔnagB (ery), ΔmanLMN (kan) | This study |
| VL6008 | D39V, prs1::PF6-tetR (gen), bgaA::Ptet-spv_0647-comEB (tet), Δspv_0647-comEB (ery) | This study |
| VL6007 | D39V, prs1::PF6-tetR (gen), bgaA::Ptet-comEB (tet), ΔcomEB (ery) | This study |
| VL6033 | D39V, prs1::PF6-tetR (gen), bgaA::Ptet-spv_0647 (tet), Δspv_0647 (ery) | This study |
| VL6477 | D39V, CEP::PF6-tetR-Ptet (spc) | Lab collection |
| VL6935 | D39V, prs1::Plac-fakB3-PF6-lacI (gen) | Lab collection |
| VL7362 | D39V, prs1::Plac-fakB3-PF6-lacI (gen), CEP::PF6-tetR-Ptet-fabM (spc), Δspv_0647 (ery) | This study |
| VL6297 | Δspv_0647 (ery) | This study |

Notes:

- Plac: IPTG-inducible promoter, PF6: constitutive promoter, Ptet: tetracycline-inducible promoter (PT5-3 variant) Sorg *et al.* (2016) [4]
- tet: tetracycline resistance, gen: gentamycin resistance, ery: erythromycin resistance, kan: kanamycin resistance, spc: spectinomycin resistance

**Supplementary Table 19. Oligonucleotides that were used in this study to construct strains.**

| ID | Sequence (5'-3') | RE* |
|---|---|---|
| OVL7886 | caactggtttaccatgcacacc | |
| OVL7887 | TCGACGTCTCGAAgagagaaagcagaagttagaga | Esp3I |
| OVL7888 | TCGACGTCTCGtcTTATTTCCTCCCGTTAAATAATAGAT | Esp3I |
| OVL7889 | TCGACGTCTCGtATGAACAAAAATATAAAATATTCTCAAAACT | Esp3I |
| OVL7890 | TCGACGTCTCGCATaatgttaacctccttaaaagattga | Esp3I |
| OVL7891 | gtattggaaccttggattgcagg | |
| OVL7892 | ggcaaagatatcgacacgg | |
| OVL7893 | CAGAGGTCTCGaaagcaatgttctattgaacgc | BsaI |
| OVL7894 | CAGAGGTCTCGcttttTTATTTCCTCCCGTTAAATAATAGAT | BsaI |
| OVL7895 | CAGAGGTCTCGagATGAACAAAAATATAAAATATTCTCAAAACT | BsaI |
| OVL7896 | CAGAGGTCTCCATcttttcatcctccatttctgtc | BsaI |
| OVL7897 | ggtccaaccagaatctgcttgg | |
| OVL8025 | ggtcacctcttgtcaagaatgc | |
| OVL8026 | GAGTCGTCTCGCATaacattttctttctacttgtcaca | Esp3I |
| OVL8027 | GAGTCGTCTCCtATGAACAAAAATATAAAATATTCTCAAAACT | Esp3I |
| OVL8028 | GAGTCGTCTCCTTTCCTCCCGTTAAATAATAGATAAC | Esp3I |
| OVL8029 | GAGTCGTCTCGGAAATAAaaattatcaaaaataaatggtttagaaagatttttaaacc | Esp3I |
| OVL8030 | ggacctgtcagcataatgatgc | |
| OVL8026 | GAGTCGTCTCGCATaacattttctttctacttgtcaca | Esp3I |
| OVL8027 | GAGTCGTCTCCtATGAACAAAAATATAAAATATTCTCAAAACT | Esp3I |
| OVL8882 | GTAGCGTCTCGttctTTATTTCCTCCCGTTAAATAATAGAT | Esp3I |
| OVL8883 | GTAGCGTCTCGagaaaaaggagaaaagagatgac | Esp3I |
| OVL8693 | caagattgctgagccacctg | |
| OVL8884 | CTCAGCTCTTCCctcttttctcctttttctctattttt | SapI |
| OVL8885 | CTCAGCTCTTCCgagATGAACAAAAATATAAAATATTCTCAAAACT | SapI |
| OVL8886 | CTCAGCTCTTCCttTTATTTCCTCCCGTTAAATAATAGAT | SapI |
| OVL8887 | CTCAGCTCTTCCAaaattatcaaaaataaatggtttagaaagatt | SapI |
| OVL3830 | CAACTCACATGAACTACATGATGAACCCAG | |
| OVL8498 | GCGTCACGTCTCAATCTTTTGAATTCGCGGCCGC | Esp3I |
| OVL5352 | GCGCTCAGCTCTTCAGATCTTTTGAATTCGCGGCCGC | SapI |
| OVL5019 | GCGTCACGTCTCAACTAGTCAAGGTCGGCAATTC | Esp3I |
| OVL5353 | GCGCTCAGCTCTTCAACTAGTCAAGGTCGGCAATTC | SapI |
| OVL5144 | AACCTGCTGCTACTGCTGCTTGGCT | |
| OVL8499 | GCGTCACGTCTCAAGATaggaggttaacattatgcctaac | Esp3I |
| OVL8500 | GCGTCACGTCTCATAGTttatgcttgataacgttttacgc | Esp3I |
| OVL8501 | GCGTCACGTCTCAAGATtggaggatgaaaagatgaaag | Esp3I |
| OVL8502 | GCGTCACGTCTCATAGTttatttttcgagtaagctaagcgc | Esp3I |
| OVL8503 | GCGTCAGCTCTTCAATCagaaagaaaatgttatgtctgaac | SapI |
| OVL8504 | GCGTCAGCTCTTCAAGTctattttttaaaaaatggtaaacc | SapI |
| OVL8879 | GCGTCACGTCTCAAGATaggagaaaagagatgactg | Esp3I |
| OVL8506 | GCGTCACGTCTCATAGTttacattagatcagcctc | Esp3I |
| OVL8880 | GCGTCAGCTCTTCAATCagaaagaaaatgttatg | SapI |
| OVL8881 | GCGTCAGCTCTTCAAGTttacattagatcagc | SapI |
| OVL10267 | Cgtcagcgttgcttggttc | |
| OVL10272 | ccaaacacctcaacaagatgg | |
| OVL10746 | GTCAcgtctcgCTGCtcgaagatttcagcttg | Esp3I |

| | | |
|---|---|---|
| OVL10747 | GTCAcgtctcgGCTTctagcaaaaaactggacg | Esp3I |
| OVL10748 | GTCAcgtctcgAAGCtcaagcaactaaaaaggaaccagg | Esp3I |
| OVL10749 | GTCAcgtctcgGCAGtattgtcattcctccttt | Esp3I |
| OVL9480 | Ggagagattcaggtcaacattgaag | |
| OVL7825 | ATGATTCTCAGACATCTGGGAATTAGC | |
| OVL10858 | CGATcacctgccgtaTAACtcgagaaaaaaaaaaccg | AarI |
| OVL10859 | CGATcacctgccgtaGTTAttaatcaatttcatagcccatcag | AarI |
| OVL10860 | CGATcacctgccgtaAATAatgacttggaagattattgctg | AarI |
| OVL10861 | GCTAcacctgccgtaTATTtttcctccttatttatttag | AarI |
| OVL10753 | Cattagccttcttatcatctcc | |
| OVL10754 | Gttgaagtcagctaagctcg | |
| OVL10678 | ATCGcgtctcgAATAatggaacacattatttatcagcttg | Esp3I |
| OVL10679 | ATCGcgtctcgATCCttattttcctataaatttaggtcttcttc | Esp3I |
| OVL4212 | GATCCGTCTCGTATTTTTCCTCCTTATTTATTTAGATCTACTCTA | Esp3I |
| OVL9724 | GATCCGTCTCGGGATCCCTCCAGTAACTCG | Esp3I |

Notes:

*RE: Restriction enzyme site introduced by PCR.
